# Cryo-EM of mammalian PA28αβ-iCP immunoproteasome reveals a distinct mechanism of proteasome activation by PA28αβ

**DOI:** 10.1101/2020.04.29.067652

**Authors:** Jinhuan Chen, Yifan Wang, Cong Xu, Chao Peng, Zhanyu Ding, Yao Cong

## Abstract

The proteasome activator PA28αβ affects MHC class-I antigen presentation by associating with immunoproteasome core particles (iCPs). However, due to the lack of a mammalian PA28αβ-iCP structure, how PA28αβ regulates proteasome remains elusive. Here we present the complete architectures of the mammalian PA28αβ-iCP immunoproteasome and free iCP at near atomic-resolution by cryo-EM, and determined the spatial arrangement between PA28αβ and iCP through XL-MS. Our structures revealed a slight leaning of PA28αβ towards the α3-α4 side of iCP, disturbing the allosteric network of the gate-keeper α2/3/4 subunits, resulting in a partial open iCP gate. We found that the binding and activation mechanism of iCP by PA28αβ is distinct from those of constitutive CP by the homoheptameric *Tb*PA26 or *Pf*PA28. Our study sheds lights on the mechanism of enzymatic activity stimulation of immunoproteasome and suggests that PA28αβ-iCP has experienced profound remodeling during evolution to achieve its current level of function in immune response.

## Introduction

Proteasomes degrade many protein substrates in the cytosol and nuclei of eukaryotic cells, and are hence essential for many aspects of cellular function (Finley, 2009; Schwartz and Ciechanover, 2009; Stadtmueller and Hill, 2011). Proteasome can refer to a variety of complexes whose enzymatic cores are the cylindrical 20S proteasome (also termed core particle, CP). Proteasome activity is fine-tuned as a result of the association of the proteolytic CP with diverse proteasomal activators (PAs), including PA700, PA28, and PA200, that can induce the opening of the CP gate into the central proteolytic cavity, allowing substrates to be degraded (Stadtmueller and Hill, 2011). PA700 (also called 19S), an ATP-dependent activator, could bind a CP to form the 26S proteasome, which mediates degradation of ubiquitylated substrates (Pickart and Cohen, 2004; Voges et al., 1999). In contrast, PA28 (also called 11S or REG in most organisms, and PA26 in *Trypanosoma brucei, Tb*PA26) stimulates the degradation of peptides in an ATP/ubiquitin-independent manner (Rechsteiner and Hill, 2005; Tanahashi et al., 2000). In the PA28 family, the homologous PA28α and PA28β, whose expressions are induced by IFN-γ, can associate into a heteroheptamer (Ahn et al., 1995; Ahn et al., 1996; Jiang and Monaco, 1997; Murata et al., 2001; Realini et al., 1994), while PA28γ can form a homoheptamer (Mao et al., 2008). PA200, also called Blm10 in yeast, is another ATP/ubiquitin-independent proteasome activator (Khor et al., 2006; Qian et al., 2013; Ustrell et al., 2002).

PA28αβ is usually linked to major histocompatibility complex (MHC) class 1 antigen processing, a critical step in immune response (McCarthy and Weinberg, 2015; Rechsteiner et al., 2000b). PA28αβ has been shown *in vitro* to affect the generation of peptides by proteasome CPs and is required for efficient presentation of many T cell epitopes from a number of viral, bacterial, and tumor-derived antigens (Rechsteiner et al., 2000a). PA28 deficiency could reduce the production of MHC class I-binding peptides in cells (de Graaf et al., 2011; Respondek et al., 2017; Schwarz et al., 2000).

Most tissues and cells express predominantly the constitutive 20S proteasome (cCP) with the proteolytic active sites located at the β1, β2, and β5 subunits of the core particle (Dick et al., 1998; Enenkel et al., 1994; Heinemeyer et al., 1997). However, lymphoid cells and cells exposed to cytokines such as IFN-γ alternatively express three homologous subunits (β1i/LMP2, β2i/MECL-1, and β5i/LMP7), replacing the constitutive ones, in the 20S immunoproteasome (iCP) particles — with this alternative expression resulting in a change in the proteolytic activities (Barton et al., 2002; Griffin et al., 1998; Khan et al., 2001; Tanaka and Kasahara, 1998; Huber et al., 2012). It has been suggested that iCPs generate class I-binding peptides to participate in antigen processing and play an important role in MHC class 1 antigen presentation (Aki et al., 1994; Hendil et al., 1998; Kloetzel, 2004).

With constant efforts during the past years, the structural basis of the 19S-cCP and Blm10-cCP systems have become better understood (Bard et al., 2018; Ding et al., 2017; Ding et al., 2019; Stadtmueller and Hill, 2011; Wehmer et al., 2017; Zhu et al., 2018). Recently, a crystal structure of mouse PA28α_4_β_3_ revealed an alternating arrangement of four α and three β chains (Huber and Groll, 2017). Still, structural studies on the PA28-CP complex are mostly limited to the homoheptameric *Tb*PA26 or *Plasmodium falciparum* (*Pf*) PA28 in complex with the cCP (Forster et al., 2005; Whitby et al., 2000; Xie et al., 2019). A complete structure of a mammalian PA28αβ-cCP or PA28αβ-iCP proteasome has not yet been determined. This deficiency is mostly due to the lability and sensitivity to salt, the binding of mammalian PA28αβ to the CP is fully reversible (Cascio, 2014). These issues make the *in vitro* assembly of the PA28αβ-CP complex or direct isolation of this complex from tissues or cells extremely challenging (Dubiel et al., 1992). As a result, the mechanism by which the heteroheptameric PA28αβ binds and activates the CP or iCP remains elusive.

Here, to examine the association and activation mechanism of iCP by PA28αβ, we assembled a mammalian PA28αβ-iCP immunoproteasome complex from separately isolated human PA28αβ and bovine spleen iCP. We determined, for the first time to the best of our knowledge, cryo-electron microscopy (cryo-EM) structures of free bovine iCP, and of the single- and double-capped mammalian PA28αβ-iCP and PA28αβ-iCP- PA28αβ complexes to resolutions of 3.3 Å, 4.1 Å, and 4.2 Å, respectively. We depicted the previously unreported spatial arrangement between the PA28αβ and the iCP by chemical cross-linking coupled mass spectrometry (XL-MS) analysis. Importantly, our structural results revealed a distinct mechanism for the binding and activation of the iCP by the heteroheptameric PA28αβ, compared with those of the cCP by the homoheptameric *Tb*PA26 or *Pf*PA28. We also found conserved differences between the immune catalytic subunits and those of the constitutive ones, beneficial for the development of immune-specific inhibitors. Our study provides insights into the unique mechanism of proteasome activation induced by PA28αβ, and how this non-ATPase activator regulates CP gate opening potentially through an on-and-off mode for substrate processing.

## Results

### Cryo-EM structures of mammalian PA28αβ-iCP immunoproteasomes

To avoid the known difficulties involved in isolating an intact PA28αβ-iCP complex directly from tissues or cells (Dubiel et al., 1992), we developed a procedure to produce a PA28αβ-iCP complex from its separately isolated components. We first expressed and purified human PA28αβ heteroheptamer from *E. coli* following the established procedure (Realini et al., 1994; Realini et al., 1997; Huber and Groll, 2017), and also isolated bovine 20S immunoproteasome directly from bovine spleen. The purifications of both proteins were confirmed using SDS-PAGE, negative-staining EM (NS-EM), and MS analyses (Fig. S1A-B). Note that we selected the spleen because this organ has been reported to display baseline immunoproteasome expression level higher than those of other organs (Ebstein et al., 2012; Noda et al., 2000). We then *in vitro* assembled the purified human PA28αβ and bovine iCP into the intact PA28αβ-iCP immunoproteasome complex in the presence of glutaraldehyde as a cross-linker (Fig. S1C-D). We further tested the *in vitro* proteolytic activity of the reconstituted PA28αβ- iCP complex against the fluorogenic peptide Suc-LLVY-AMC; the results of this test showed that the complex was functionally active (Fig. S1E).

From the same set of cryo-EM data of the mammalian PA28αβ-iCP complex, we resolved three maps, including the free bovine iCP, the single-capped PA28αβ-iCP and the double-capped PA28αβ-iCP-PA28αβ at the resolutions of 3.3 Å, 4.1 Å, and 4.2 Å, respectively (Figs. 1A-D, S2, and Table S1). To the best of our knowledge, none of these structures has been determined previously. Our PA28αβ-iCP and PA28αβ-iCP- PA28αβ maps revealed a funnel-like PA28αβ heteroheptamer associated with, respectively, one or both ends of the iCP (Fig. 1A-B). PA28αβ was observed to be ∼90 Å in diameter and ∼90 Å in height, and to consist of a central channel 35 Å in diameter at the 20S-binding end and 20 Å at the distal end (Fig. 1A). The size of the channel was found to be comparable to those of the homologous PA26 and PA28 systems (Huber and Groll, 2017; Xie et al., 2019). Note that our cryo-EM maps revealed extra pieces of density extending on top of the PA28αβ core, to some extent covering the entrance to the funnel; these pieces of density most likely derived from the dynamic apical loops (Fig. 1A-B). Additional local resolution analysis suggested that in both capped complexes, PA28αβ appeared less well resolved than did the complexed iCP, indicating the intrinsic dynamic nature of PA28αβ, especially in its unstructured apical loop region (Fig. 1E-F). This dynamic nature of the apical loops may potentially be beneficial for substrate recruitment and substrate entry into or efflux out of the central channel of PA28αβ.

**Fig. 1.**
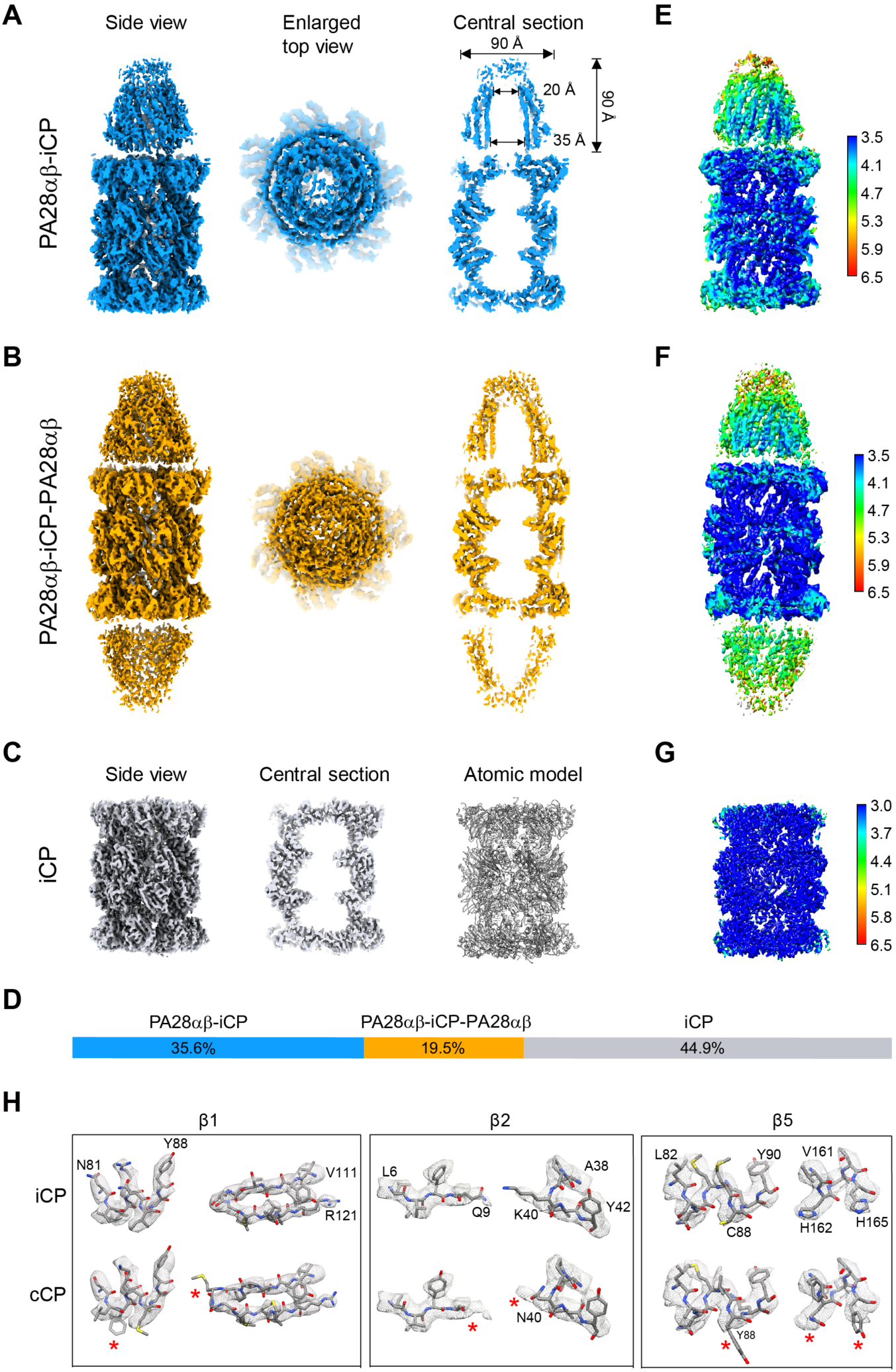
Overall cryo-EM maps of mammalian PA28αβ-iCP, PA28αβ-iCP-PA28αβ and iCP. **(A)** Cryo-EM map of the single-capped PA28αβ-iCP complex (side view, enlarged top view and one central section). **(B)** Cryo-EM map of the double-capped PA28αβ-iCP-PA28αβ complex, in the same rendering style as in **(A). (C)** Cryo-EM structure of the free iCP reconstructed from the same dataset (side view, one central section, and atomic model). **(D)** The relative populations of PA28αβ-iCP (dodger blue), PA28αβ-iCP-PA28αβ (gold), and iCP (gray). **(E-G)** Local resolutions of the PA28αβ- iCP **(E)**, PA28αβ-iCP-PA28αβ **(F)**, and iCP **(G)** cryo-EM maps determined using ResMap. Shown are the cut-away view of the corresponding density map, with the color bar on the right labeling the resolutions (in Å). **(H)** Identifications of immunoproteasome amino-acid residues by fitting details. The iCP sequence (top row), but not the conventional CP sequence (bottom row), fits well into the density of our iCP cryo-EM map in the three catalytic subunits. The red asterisks indicate the residues of cCP that do not match well with the density.

For the free iCP, a considerable portion of the map showed local resolution levels better than 3.0 Å (Fig. 1G), revealing side chain densities of most of the amino acid residues, and allowing us to build a pseudo-atomic model of the complex (Fig. 1C). Note that the unambiguous assignment of amino acid side chains here enabled us to confirm that our purified bovine spleen 20S proteasome was indeed the immunoproteasome, with their constitutive counterparts substituted from β1, β2, and β5 to β1i, β2i, and β5i, respectively (Fig. 1H).

### Relative spatial arrangement between PA28αβ and iCP determined by XL-MS

A recent crystal structure of mouse PA28α_4_β_3_ revealed an alternating arrangement of four α and three β chains with two consecutive α subunits sitting side-by-side (Huber and Groll, 2017). Due to the high sequence identity (∼94%) between human and mouse PA28αβ (Fig. S3A) and the similar expression and purification procedure undertaken by us and in their study (Huber and Groll, 2017), the subunit order of human PA28αβ heteroheptamer was expected to be the same to that of mouse PA28αβ (Huber and Groll, 2017), which is the case for the subunit ordering of the more complexed eukaryotic chaperonin TRiC/CCT consisting of eight paralogous subunits (Kalisman et al., 2013; Leitner et al., 2012; Zang et al., 2016; Zang et al., 2018). Still, the relative spatial arrangement of the PA28αβ and CP units remained unclear. We therefore carried out an XL-MS analysis of PA28αβ-iCP to determine this arrangement. This analysis disclosed a number of interactions involving the C-termini of the PA28α or PA28β subunit and the amino acid residues lying in the iCP α-ring pocket regions, including PA28α-α2, PA28α-α3, PA28β-α2, and PA28β-α6 interactions (Figs. 2A and S3B). These interaction constraints led to the deciphering of a unique spatial arrangement of the PA28αβ and iCP units relative to each other, with the two consecutive PA28α subunits (α_1_ and α_4_) residing on top of the α6 and α7 subunits of iCP (Fig. 2B). This was the first determination of the relative spatial arrangement between PA28αβ and iCP.

**Fig. 2.**
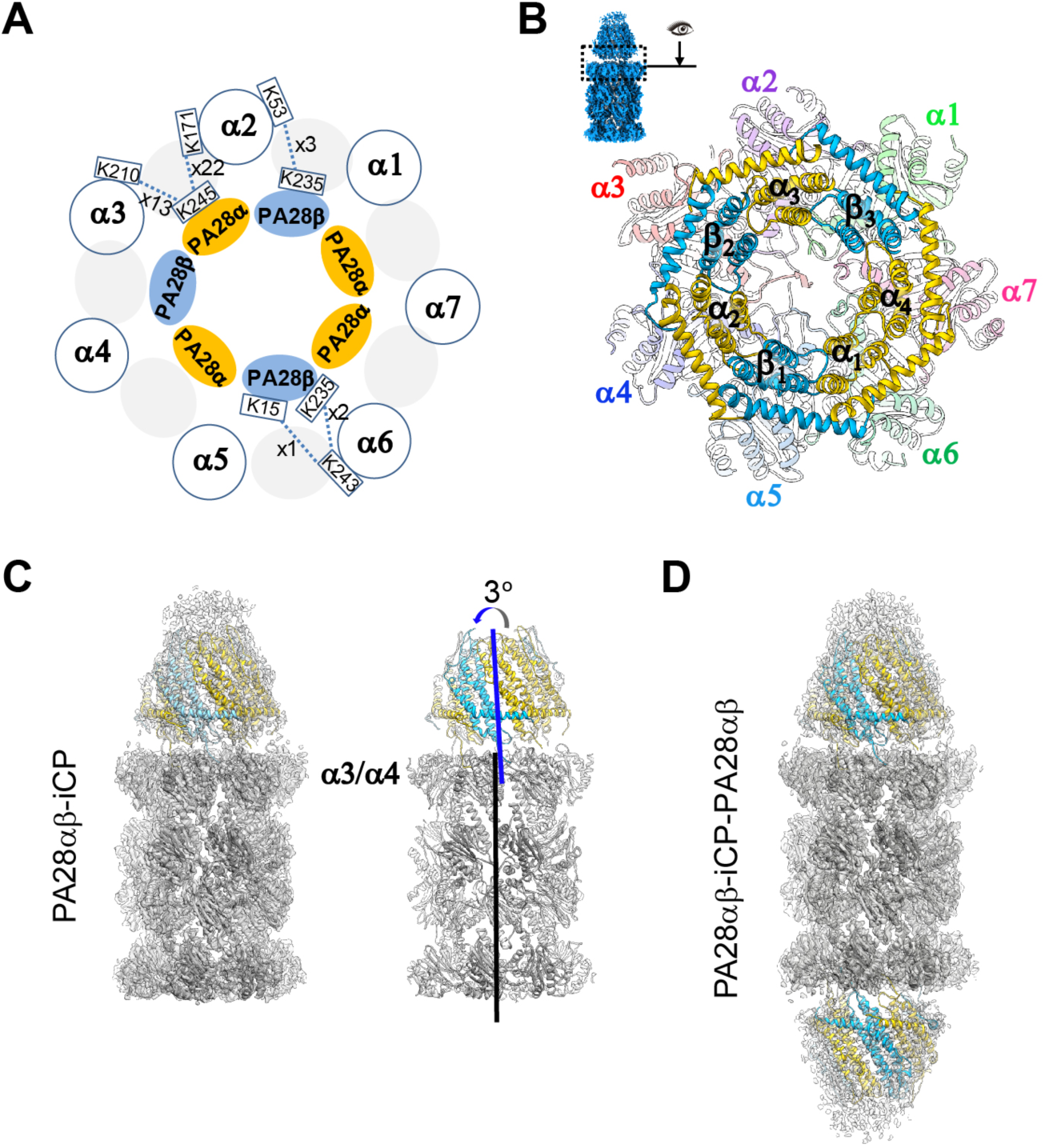
The spatial arrangement between PA28αβ and iCP in the PA28αβ-iCP proteasome. **(A)** A carton diagram illustrating the spatial arrangement between PA28αβ and iCP (blue circles) derived from XL-MS analysis. The PA28α and PA28β are shown as sky blue and gold ellipses, respectively, and the color scheme was followed throughout. Identified cross-linked contacts (only in the interface) between a pair of subunits are shown as dotted lines, with the involved residues and the spectrum count number indicated. We used an E-value of 1.00E^-02^ as the threshold to remove extra lower-confidence XL-MS data. **(B)** Visualization of the PA28αβ-iCP interface, with the iCP subunits shown in distinct colors and appearing transparent. The visualization angle and region are illustrated in the Inset. **(C)** Cryo-EM map of PA28αβ-iCP (transparent grey) with fitted model (iCP in grey, PA28αβ in color, left panel). On the right showing the model of PA28αβ-iCP, with the position of the α3/α4-subunit and the axes of PA28αβ (blue) and iCP (black) indicated. The axes also indicate the direction of the tilt of PA28αβ relative to iCP. **(D)** Cryo-EM map and model of PA28αβ-iCP-PA28αβ.

Based on this relative spatial arrangement of the PA28αβ and iCP units, we built a pseudo-atomic model for each of the PA28αβ-capped complexes (Fig. 2C-D). Interestingly, our structure revealed a slight tilt (∼3°) of PA28αβ toward the α3-α4 side of the iCP (Fig. 2C), comparable to that observed for the *Pf*PA28-CP complex (Xie et al., 2019). This lack of alignment of the axes of the PA28αβ and iCP units was reminiscent of a resting state 26S proteasome with the Rpt ring lean sitting on the plane of the 20S CP (Ding et al., 2019).

### A unique mode of interaction between the mammalian PA28αβ activator and iCP

Based on our PA28αβ-iCP structure, we analyzed the interaction interface between PA28αβ and the iCP. The activator C-terminal tail insertions into the CP pockets have been documented to be able to stabilize the binding of activator but to not be sufficient to open the CP gate (Smith et al., 2007; Whitby et al., 2000). Our PA28αβ- iCP cryo-EM map showed relatively strong pieces of density corresponding to four PA28αβ C-terminal tail insertions into the α-ring pockets of the iCP, including the C-termini of PA28 β_3_ α_3_, β_2_, and α_2_ inserted into, respectively, the pockets of α1/2, α2/3, α3/4, and α4/5 of the iCP, and a rather weak piece of density for the C-terminus of PA28β_1_ in the α5/6 pocket (Fig. 3A-B). However, we found no obvious extra density in the pockets of α6/7 and α7/1 (Fig. 3A-C). Interestingly, PA28αβ overall displayed stronger interactions with the α3-α4 side of the iCP but no binding with the opposite α6-α7 side, consistent with our above-described observation of the slight leaning of PA28αβ towards the α3-α4 side, and resulting formation of more intimate interactions with the iCP on this side. Surprisingly, the iCP α-ring pocket occupancy status in our mammalian PA28αβ-iCP differs considerably from that described for *Pf*PA28-cCP, which showed only one weak *Pf*PA28 C-terminal insertion into the α7/1 pocket of its cCP although it showed a slight tilt of the activator toward α3-α4 (Fig. 3C) (Xie et al., 2019). In contrast, the occupancy status in our PA28αβ- iCP differs only a little from that for *Tb*PA26-cCP, with the only major difference being a lack of a PA26 C-terminus insertion into the α1/2 pocket of the cCP (Fig. 3C) (Forster et al., 2005). Taken together, our data suggested the binding mode and mechanism of association between the heteroheptameric PA28αβ and the iCP (and most likely also for the cCP) to be quite different from that described for the homoheptameric *Pf*PA28 with the cCP in terms of strength and location, but to be more comparable to that of *Tb*PA26 with the cCP.

**Fig. 3.**
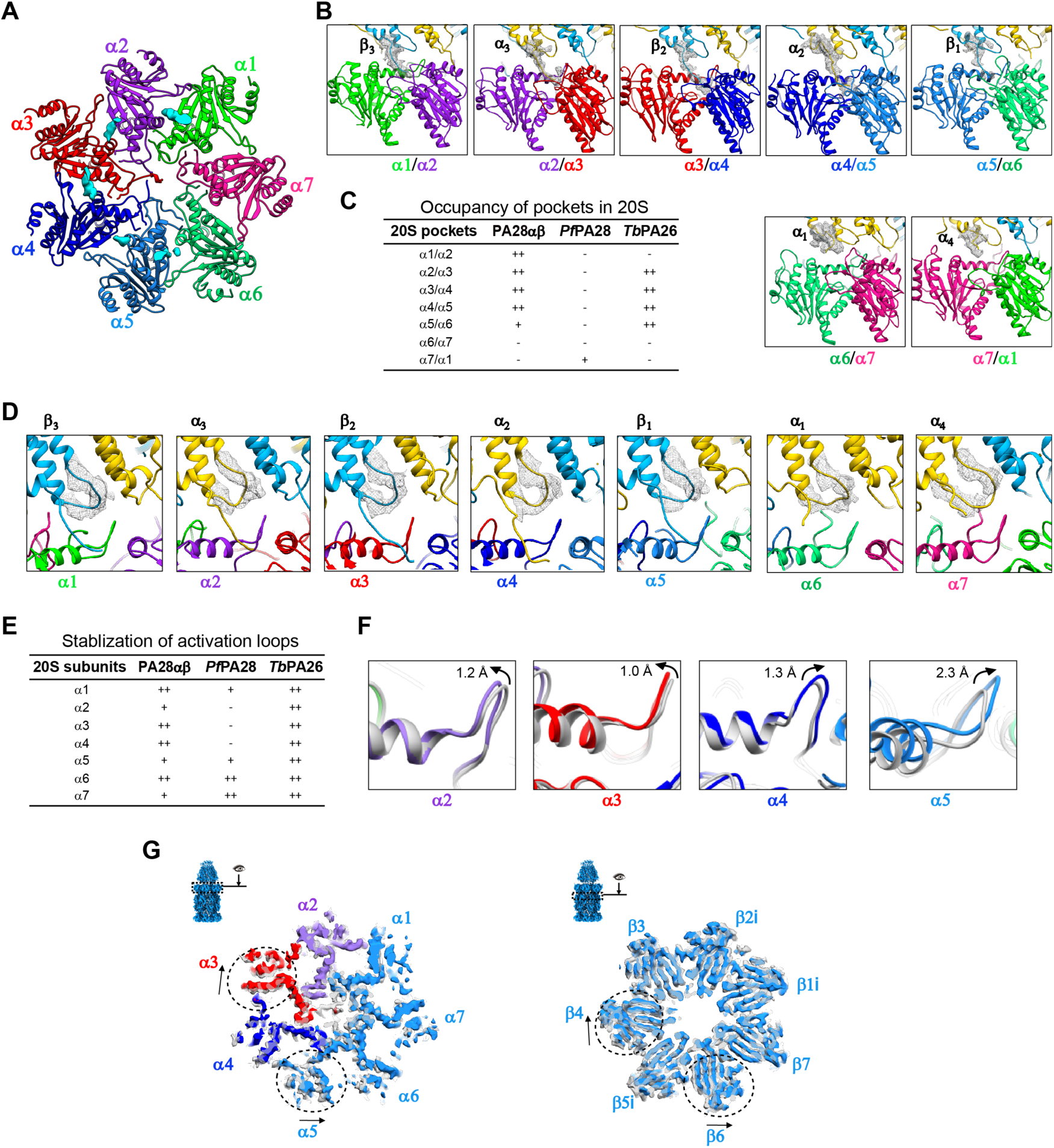
Major interactions between PA28αβ and iCP and the induced conformational changes of iCP. **(A)** Top view of the iCP α-ring with the inserted C- terminal tails of PA28αβ (cyan pieces of density) for the PA28αβ-iCP complex. **(B)** Magnified views of the insertions of PA28αβ C-termini into the iCP α-ring pockets, with the C-tail density shown as grey wire. The color schemes for the model of PA28αβ and iCP are as in Fig. 2B, and are followed throughout. **(C)** Occupancy status of the 20S pockets of PA28αβ, *Pf*PA28 and *Tb*PA26. ++: strong density, +: weak density; -: no density. **(D)** The activation loop status of PA28αβ (the loop density shown as grey wire). **(E)** Stabilization status of activation loops of PA28αβ, *Pf*PA28, and *Tb*PA26. ++: well resolved, +: less well resolved; -: not resolved. **(F)** Reverser turn conformational changes induced by PA28αβ association, determined by aligning the models of PA28αβ-iCP (in color) with that of the free iCP (grey). Black arrow indicates the shift direction. **(G)** Superpositions of the cis α-rings (left) and β-rings (right) of PA28αβ-iCP (with color) and the free iCP (grey), showing their conformation differences. Dashed circles indicate regions with considerable conformational changes between the two structures, and arrows show the direction of conformational switches. The visualization angle and region are illustrated in the Inset.

Furthermore, another critical contact between PA26 and the cCP has been indicated to involve the formation of an interaction between the activation loop of PA26 and the reverse turn of the CP α-subunit (located between the N-terminal tail and H0 of α-subunit), and in part lead to the gate opening of the CP (Whitby et al., 2000). Indeed, the *Tb*PA26-cCP structure shows interactions between all of the activation loops of the homoheptameric PA26 and the related reverse turns of the CP (Whitby et al., 2000). In our PA28αβ-iCP structure, all seven activation loops of PA28αβ were resolved, with the ones interacting with α1, α3, α4 and α6 of iCP showing stronger density (Fig. 3D-E). In contrast, the activation loop-reverse turn interaction in the *Pf*PA28-cCP structure was described to be very different, and showed interactions only on the α6-α7 side of the CP (including α5, α6, α7, and α1), with the interactions with α6 and α7 appearing stronger, but did not reveal the activation loop densities interacting with the gate keeper α2, α3, and α4 subunits (Fig. 3E) (Xie et al., 2019).

In addition, we also observed conformational changes in the iCP induced by PA28αβ binding, including a slight shift (up to 2.3 Å) in the reverse turn regions of the gate keepers α2/3/4 and the neighboring α5 in the iCP (Fig. 3F). In addition, we also found slight rotations of the peripheral portions of α3 and α5 surrounding the α4 subunit; these rotations could play a role in disturbing the allosteric network of the gate keepers α2/3/4 (and to a lesser extent of α5) (Fig. 3G) These motions may have been induced by the noticeable leaning of PA28αβ towards the α3-α4 side of the iCP, and could be propagated to the β-ring with visible movements of the underneath β4 and β6 subunits (Fig. 3G). In contrast, *Pf*PA28 binding was indicated to not induce any large changes in the conformations of the α-ring subunits of the cCP (Xie et al., 2019). While for the *Tb*PA26-cCP proteasome complex, *Tb*PA26 binding was indicated to result in a rearrangement of the N-terminal extensions of α2, α3, α4 and α5 to a conformation similar to that of α6, α7, and α1 and in this way to an opening of the axial pore (Whitby et al., 2000).

Taken together, our data indicated that, relative to the homoheptameric *Pf*PA28 or *Tb*PA26, the heteroheptameric mammalian PA28αβ interacts with and activates the enzymatic core particle using related but different mechanisms, especially distinct from that of *Pf*PA28; consequently, the motions they induced to CP are also divergent. These observations, along with the relative low sequence identities between these proteins from different species (Fig. S4), suggested that during evolution the heteroheptameric PA28αβ has undergone profound remodeling to achieve its current physiological functions including immune response in eukaryotes.

### The mechanism of PA28αβ-induced partial gate opening of the iCP

Activation of proteasomes largely relies on the gate opening of the CP, which could be triggered by regulators or together with substrates (Ding et al., 2017; Forster et al., 2005; Sadre-Bazzaz et al., 2010; Whitby et al., 2000). Our cryo-EM map of the free iCP showed a closed-gate configuration with ordered and well-resolved N-termini of the α2, α3, α4, and to a lesser extent α5 subunits showing extended conformations covering the gate region (Fig. 4A); while the N-termini of the remaining α6, α7, and α1 subunits were observed to point away from the pore and to approximately align with the central axis of the iCP, thus not participating in blocking the iCP pore. In contrast, in our PA28αβ-iCP map, the N-termini of the α2, α3, and α4 subunits in the PA28αβ- contacting iCP α-ring appeared disordered, suggesting that the binding of the PA28αβ activator to iCP caused these regions to become more dynamic (Fig. 4B). This plasticity in these N-terminal regions may have disrupted the allosteric networks in the gate region, leading to a partially open gate in the contacting α-ring, while leaving the gate in the opposite α-ring closed (Fig. 4B). Consistent with this proposal, both gates appeared to have adopted a partially open conformation in our double-capped PA28αβ- iCP-PA28αβ map (Fig. 4C). Thus, the binding of the PA28αβ activator to the iCP was concluded to have induced a partial gate opening of the iCP.

**Fig. 4.**
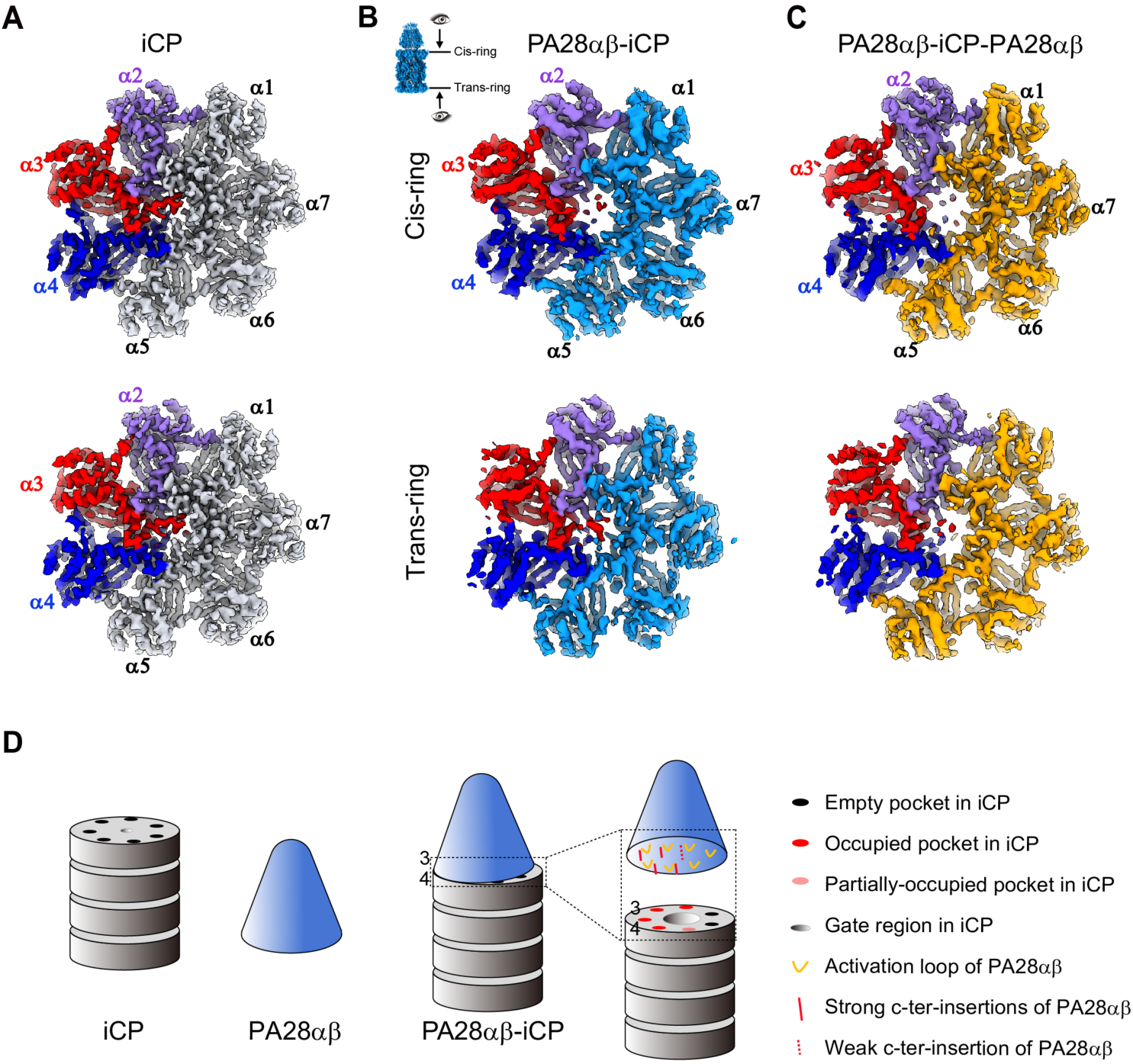
The states of iCP gates in bovine immunoproteasomes and a proposed gate opening mechanism induced by PA28αβ. **(A)** The gate status of the free bovine iCP, with both rings showing a closed-gate conformation. The gate keepers α2, α3, and α4 are shown in purple, red and blue, respectively, which scheme is followed throughout this figure. **(B)** The gate status of bovine PA28αβ-iCP, with the PA28αβ-capped cis-ring showing a partially open gate, while the trans-ring still showing a closed gate. The visualization angle and region are illustrated in the Inset. **(C)** The gate status of PA28αβ- iCP-PA28αβ, with both the cis-ring and trans-ring showing a partially open gate. **(D)** A schematic diagram depicting a proposed mechanism of gate opening induced by PA28αβ binding.

In general, there are two important elements of proteasome activation by 11S activators: the C-termini of the activator, which provide binding energy; and the activation loop–reverse turn interaction that can destabilize the closed-gate conformation (Whitby et al., 2000). Here, by comparing the structure of the activator-associated PA28αβ-iCP complex with that of the free iCP, we derived a detailed proposed mechanism for the iCP partial gate opening induced by the mammalian PA28αβ activator (Fig. 4D). According to this mechanism, the association of PA28αβ with the iCP would result in the insertion of the C-terminal tails of four consecutive PA28αβ subunits (β_3_, α_3_, β_2_, and α_2_) into the corresponding α-pockets (α1/2, α2/3, α3/4, and α4/5), causing PA28αβ to lean toward the α3-α4 side of the iCP. The close proximity in this position could facilitate the formation of interactions between the activation loops of PA28αβ and the reverse turns of the iCP subunits, especially the α2/3/4 gate keepers, resulting in a shift in their reverse turn regions. As the N-termini of α1, α6, and α7 have been observed to adopt conformations pointing away from the proteasome, they would contribute less to the gate formation, while α2/3/4, and to a lesser extent α5, have been observed to pack closely to cover the gate. Thus, the interactions between the reverse turns of α2/3/4 and the corresponding PA28αβ activation loops would disrupt the allosteric networks of the gate keepers, resulting in a partially open gate in PA28αβ-iCP.

### Unique properties of the beta catalytic subunits in bovine immunoproteasomes

During infection of antigens, PA28αβ as well as the β1i, β2i, and β5i subunits of the core particle would be induced by INF-γ to form immunoproteasomes, facilitating the generation of MHC class 1 ligands for subsequent antigen presentation (Griffin et al., 1998; Khan et al., 2001; Rivett and Hearn, 2004). However, the underlying molecular mechanism responsible for the stimulation of immunoproteasome activity remains unclear. Here we first asked whether the association of PA28αβ could trigger considerable conformational changes of the three catalytic subunits that may play a role in stimulating the immunoproteasome activity. Our subunit alignment analysis suggested that the association of PA28αβ with iCP would tend to have a subtle effect on the conformations of β1i and β2i (Fig. S5A), although it could slightly reshape a turn located outside the chamber and helix 3 (H3) of β5i (located in the outermost region of the chamber) (Fig. S5A). This analysis suggested that the production of antigen ligands in the iCP may mostly arise from the replacement of standard beta subunits with catalytic subunits.

We then compared the conformations of the three catalytic subunits of our free bovine iCP structure with those of the available bovine cCP structure (PDB ID: 1IRU) (Unno et al., 2002). This comparison showed noticeable conformational differences between them in several regions (Fig. 5A). For instance, relative to the cCP, the C-terminal tail of β1i in the iCP showed a distinct conformation and a linker region (Gly133-Leu139) with a noticeably rearranged structure, the C-terminal loop of β2i exhibited a slight outward displacement, and the β5i H3 displayed an observable outward shift (Fig. 5A). Interestingly, we observed similar conformational differences between the human iCP and cCP and between the mouse iCP and cCP (Fig. S5B-C). These data suggested that the conserved conformational changes and related elements may play a role in immunoproteasome activity stimulation. Furthermore, for the bovine proteasome, the patterns of surface property in the catalytic pockets of β1i were observed to be different from those of β1 (Fig. 5B), with this difference mainly resulting from the difference between β1-Thr31 and β1i-Phe31. Interestingly, a similar amino acid residue and corresponding surface property difference was observed between human β1i and β1, as well as between mouse β1i and β1 (Fig. S5D), indicating these conserved differences may contribute to the suppression of caspase-like activity in β1i in mammalian iCPs (Ferrington and Gregerson, 2012). Taken together, these findings may to some extent facilitate our understanding of the mechanism of immunoproteasome activity stimulation.

**Fig. 5.**
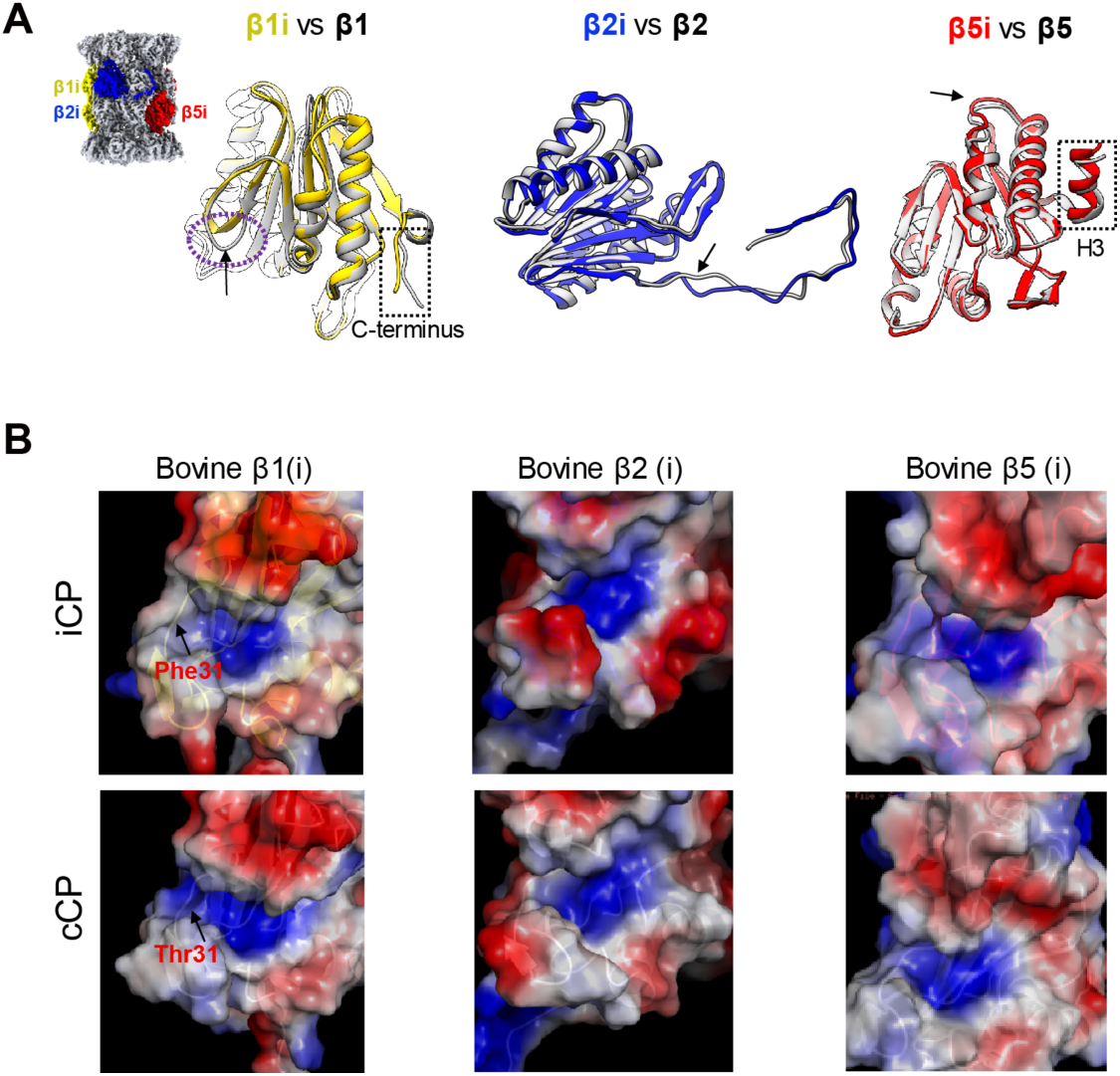
Structural comparison of the free bovine iCP with the constitutive bovine CP. **(A)** Structural superpositions of the three catalytic subunits of the free bovine iCP (with β1i, β2i, and β5i colored yellow, blue and red, respectively) on the corresponding subunits of the bovine cCP (in gray, PDB ID: 1IRU) (Unno et al., 2002). The observable conformational changes are indicated by black arrow and dotted ellipsoid or rectangle, which are followed throughout. **(B)** Surface charge representations of the catalytic pockets of the three enzymatic subunits for free bovine iCP and bovine cCP, with the most distinct residues between β1i and β1 in this region indicated.

## Discussion

Unlike 19S activator, the 11S activators (PA28/REG and PA26) stimulate the degradation of substrates in an ATP/ubiquitin-independent manner. Within the 11S family, PA28αβ forms a heteroheptamer while the remaining members (PA28γ and PA26) form respective homoheptamers. PA28αβ plays an important role in MHC class I antigen presentation (Raule et al., 2014; Tanaka and Kasahara, 1998). To date, there remains a lack of structural information on the intact mammalian PA28αβ-cCP or the PA28αβ-iCP complex, and thus the mechanism by which PA28αβ binds and activates iCP is elusive. Here we provide insight into these mechanisms by acquiring and analyzing the first cryo-EM structure of mammalian PA28αβ-iCP immunoproteasome complex with degradation ability as well as the structure of free bovine spleen iCP. We determined the spatial arrangement of PA28αβ and iCP units relative to each other, which allowed us to derive a plausible mechanism for the interaction and activation of iCP by the heteroheptameric PA28αβ, largely distinct from the mechanisms previously proposed for cCP by the homoheptameric *Tb*PA26 or *Pf*PA28. Our findings suggested that PA28αβ-iCP has experienced a profound remodeling during evolution to achieve its current level of function in immune response.

### A distinct proteasome activation mechanism adopted by mammalian PA28αβ

Although human PA28αβ, *Pf*PA28, and *Tb*PA26 are all 11S activators, they appear to cause CP gates to be opened to different extents. In contrast to the fully open gate of the CP in the *Tb*PA26-CP proteasome complex, here our structural study demonstrated that human PA28αβ binding could cause the formation of a partially open gate of the iCP (Fig. 4). This difference in the CP gate opening status induced by the two proteasome activators could be attributed to several factors: (1) their different oligomerization states, with human PA28αβ being heteroheptameric but *Tb*PA26 homoheptameric; (2) their different orientations relative to the CP, with PA28αβ being slightly tilted, but *Tb*PA26 untilted; and (3) their different interactions with the CP, with the binding of human PA28αβ to iCP perhaps being overall stronger (with one more C-terminal insertion), but the activation loop–reverse turn interactions of *Tb*PA26 appearing more evenly distributed and stronger (Fig. 3C and E) which could facilitate the full opening of the CP gate.

Moreover, although *Pf*PA28-CP also in the previous investigation showed a slight leaning of the activator to the α3-α4 side and a partially open CP gate, similar to that observed between PA28αβ and the iCP, the mode of interaction between *Pf*PA28 and CP appeared very different, with this interaction obviously weaker than that of PA28αβ with the iCP. For instance, human PA28αβ showed five C-terminal tail insertions into the pockets of the α-subunits around the α3-α4 side, while *Pf*PA28 displayed much weaker interactions with only one insertion into the opposite side (the α7/1 pocket) (Fig. 3B-C). Moreover, our PA28αβ-iCP structure allowed us to capture all seven activation loops stabilized by the interaction with the reverse turn of the α-subunits (albeit with three of them being relatively dynamic) (Fig. 3D-E). The *Pf*PA28-cCP structure showed four such interactions only on the α6-α7 side of the CP (with two of them appearing stronger), and no pieces of density were observed corresponding to activation loops interacting with the α2, α3, and α4 gate keepers (Fig. 3E) (Xie et al., 2019). Taken together, the above analysis implied that the mammalian PA28αβ may use a mechanism distinct from the mechanisms used by *Pf*PA28 and *Tb*PA26 to activate the enzymatic core particle.

### A potential on-and-off mode to regulate the CP gate opening by PA28αβ

Compared with the non-ATPase PA28αβ activator, the interaction and activation mechanism exercised by the AAA+ ATPase 19S activator has been indicated to be more precisely regulated and to involve different structural elements, with the C-terminal tails of several Rpt subunits from the hexameric ATPase ring inserted into the related α-ring pockets of CP, without direct interactions formed by activation loops and reverse turns as in the case of PA28αβ. Also, the number of inserted Rpt C-terminal tails has been indicated to vary during its ATP cycle, with an extra Rpt6 or Rpt6/1 C-tail insertion (into the pockets in the α2/3/4 side) used to trigger the opening of the 20S gate (Chen et al., 2016; Ding et al., 2019; Wehmer et al., 2017). In contrast, for PA28αβ-iCP, we showed that the slight leaning of the PA28αβ relative to the iCP could lead to a more intimate interaction with the α2/3/4 subunits (also serving as the gate keepers) of the core particle, facilitating the gate opening of the CP and forming an active state. Interestingly, although the activation seems to occur through different structural elements, they all regulate the allosteric network of the same enzymatic machine, i.e., the 20S CP, and eventually disturb the allosteric network in the key α2/3/4 gate keepers to trigger the gate opening, readying the proteasome for substrate translocation and processing within the enzymatic core. This suggests that the 20S CP has evolved to be complex enough that uses one set of allosteric network to take signals from diversified activators; still, the sophisticated machine can eventually collect all the signals and converge them to touch the ultimate key trigger point to activate the entire assembly.

Furthermore, for the ATPase-containing 19S activator, binding is not sufficient to trigger the CP gate opening; instead, it requires ATP to eventually drive the CP gate status varying between closed and open facilitating substrate processing (Wehmer et al., 2017; Zhu et al., 2018). In contrast, for the non-ATPase PA28αβ activator, it appeared that the binding and activation could happen simultaneously, i.e., once the PA28αβ activator associates with the CP, its gate would open (Fig. 4B-C). Also given the difficulty of a steady association between PA28αβ and CP observed here as well as in a previous study (Dubiel et al., 1992), we postulated that the non-ATPase PA28αβ activator likely uses an on-and-off mode (association and disassociation with the CP) to regulate CP gate opening and closing and substrate processing, as may also be the case for other non-ATPase proteasome activators. This may reflect that, during the course of evolution, the physiological role of the activator and the divergent level of the substrates determine the structural complexity of the activator and the mechanism of CP gate regulation.

### Mechanism of enzymatic activity stimulation of bovine immunoproteasome

The formation of immunoproteasomes induced by INF-γ could enhance proteasomal trypsin-like (β2) and chymotrypsin-like (β5) activities and suppress caspase-like activity (β1) (Driscoll et al., 1993; Gaczynska et al., 1993; Groettrup et al., 1995), facilitating the generation of MHC class 1 ligands for subsequent antigen presentation. In addition, INF-γ-induced expression of PA28αβ could markedly stimulate proteasomal degradation of short peptides *in vitro* (Dubiel et al., 1992; Ma et al., 1992), leading to substantial changes in the pattern of peptides generated (Raule et al., 2014). However, due to the absence of a PA28αβ-iCP structure, little is known about the molecular mechanism responsible for the immunoproteasome activity stimulations.

Here we found some noticeable conserved conformational differences in several loops of β1i and β2i, and the H3 helix of β5i between the iCP and cCP in bovine, human, and mouse (Figs. 5 and S5), and these differences might play a role in immunoproteasome functions. Also, we observed a surface property difference in the catalytic center of β1i distinct from that of β1 (Fig. 5B), mainly resulting from the primary sequence variation between β1i and β1 in the enzymatic pocket. Interestingly, this phenomenon was found to be conserved among human, mouse, and bovine (Fig. S5), which could contribute to the lower caspase-like (β1) activity of the immunoproteasome (Driscoll et al., 1993; Gaczynska et al., 1993; Groettrup et al., 1995). Taken together, the variant amino acid residues and resulting distinct surface properties may contribute to the regulation of iCP immunoproteasome activity, especially for β1i. Furthermore, this finding may be beneficial for the development of immunoproteasome inhibitors, especially β1i-specific inhibitors.

### Potential effect of PA28αβ on substrate processing

Substrates have been proposed to go through the central channel running along PA28αβ to enter the chamber of the CP (Sugiyama et al., 2013). Also, PA28αβ may control the efflux of longer peptides out of the proteolytic chamber and contribute to their ongoing hydrolysis. Hence PA28αβ could serve as a selective sieve that controls the entry of substrate and/or the exit of degradation products (Xie et al., 2019). The quite narrow channel entrance (20 Å) of PA28αβ may, to some extent, impose a stretching or unfolding force on the engaged substrates/products to regulate their entry/exit (Fig. 1A). Besides, our structure also allowed us to visualize the substrate-recruitment loops of PA28αβ, which tend to form a dome loosely covering the entrance of the PA28αβ channel (Fig. 1A-B). These loop regions show relatively lower local resolution, indicating an intrinsic dynamic nature of these regions. The plasticity may be beneficial for these PA28αβ loops to be involved in the substrate recruitment and selection at the entrance of the complex, and could even be displaced during the entrance/translocation of substrates into the channel of PA28αβ for subsequent degradation or exit of the intermediate/final product.

In summary, our study has revealed the first complete architecture of mammalian immunoproteasome PA28αβ-iCP and that of the bovine iCP, and has provided new insights into a distinct mechanism by which PA28αβ might activate the proteasome. Our data have also partially clarified the regulation mechanism of iCP immunoproteasome activity, especially for β1i, beneficial for inhibitor development. Interestingly, we delineated a mechanism on how the sophisticated 20S CP has evolved to use one set of allosteric network to take signals from diversified activators, and eventually converge these signals to touch the ultimate key trigger point to open the gate. We also proposed an on-and-off mode by which the non-ATPase activator such as PA28αβ likely uses to regulate CP gate opening and closing, and provided insights in the potential effect of PA28αβ on substrate processing.

## Materials and methods

### Molecular biology and purification of PA28αβ

For co-expression of human PA28α and PA28β, their DNA fragments were inserted into the pETDuet-1 vector system (MSC1: N-terminal-6xHis-PA28α; MSC2: N-terminal-Flag- PA28β). Reconstructed plasmids were transformed into *E. coli* BL21 (DE3) cells and selected for resistance to ampicillin on agar plates. Transformants were grown at 37°C in liquid LB medium supplemented with ampicillin. Gene expression was induced using 1 mM IPTG. After induction for 16 hours (h) at 18°C, cells were harvested and frozen at -80°C. The frozen *E. coli* cells were lysed with an ultra-high-pressure cell disrupter in lysis buffer (50 mM Tris-HCl, pH 7.5, 100 mM NaCl, 1 mM DTT, 10% glycerol). The lysate was centrifuged at 20,000 g for 30 min at 4°C. The clarified lysate was incubated with Ni-NTA agarose beads (Sigma) for 30 min at 4°C. The beads were recovered and washed twice with wash buffer (50 mM Tris-HCl, pH 7.5, 100 mM NaCl, 1 mM DTT, 10% glycerol, 20 mM imidazole) before eluting with

50 mM Tris-HCl, pH 7.5, 100 mM NaCl, 10% glycerol, 200 mM imidazole. The collected elution was subsequently incubated with anti-FLAG M2 agarose beads (Sigma) for 2 h at 4°C. Then, the beads were recovered and washed twice with wash buffer (50 mM Tris-HCl, pH 7.5, 100 mM NaCl, 10% glycerol) before eluting with 500 µg/ml 3×FLAG peptide (Shanghai Biotech BioScience & Technology). The resulting samples were concentrated by using 100 kDa centrifugal filter devices and subjected to size-exclusion chromatography (Superdex 200 16/60; GE Healthcare) with 20 mM Tris-HCl, pH 7.5, 100 mM NaCl, and 1 mM DTT. The fractions were further analyzed using SDS-PAGE (Fig. S1), western blot and MS (Fig. S1). For long-term storage, the PA28αβ samples were supplemented with 15% glycerol and stored at -80°C.

### Purification of bovine iCPs

Bovine iCPs were purified from bovine spleen following a procedure similar to that described previously (Leggett et al., 2005). Bovine spleen (50 g) was cut into pieces and homogenized in lysis buffer (25 mM Tris-HCl (pH 7.4), 10 mM MgCl_2_, 4 mM ATP, 1 mM DTT, 10% glycerol). The homogenate was centrifuged at 20,000 g for 30 min and then centrifuged at 100,000 g for 1 h to remove cell debris and membranes. The supernatant was applied to a 100 mL DEAE-Affigel Blue (Bio-Rad) column equilibrated with buffer A (25 mM Tris-HCl (pH 7.4), 10 mM MgCl_2_, 1 mM ATP, 1 mM DTT, 10% glycerol). Then the resin was washed with buffer A and 50 mM NaCl in buffer A. Proteasomes were eluted with buffer AN (25 mM Tris-HCl (pH 7.4), 10 mM MgCl_2_, 1 mM ATP, 1 mM DTT, 10% glycerol) and directly applied to a 50 mL Source 15Q column equilibrated with buffer AN. The source 15Q column was washed with buffer AN and eluted with a 500 mL gradient of 150-500 mM NaCl in buffer A. Proteasomes were eluted at a salt concentration of 300-330 mM NaCl and the activities of the collected fractions were monitored by performing a peptidase activity assay (described below). Fractions containing proteasomes were desalted and concentrated, and finally purified by using a glycerol gradient (15-45% glycerol (wt/vol), 25 mM Tris-HCl (pH 7.4), 10 mM MgCl_2_, 1 mM DTT) and subjecting them to centrifugation for 16 h at 37,000 rpm. Note that the fractions containing iCP and 26S were identified by performing SDS-PAGE and negative-staining EM as well as the peptidase activity assay (Fig. S1). The samples were frozen using liquid nitrogen and stored at -80 °C.

### In vitro reconstitution

Before mixing together the above-purified PA28αβ and iCP products, the PA28αβ product was dialyzed against buffer1 (20 mM Hepes, pH 7.5, 100 mM NaCl, 10% glycerol, and 1 mM DTT), and the iCP product was dialyzed against buffer2 (25 mM Hepes (pH 7.4), 10 mM MgCl_2_, 1 mM DTT, 10% glycerol). The dialyzed PA28αβ was mixed with the dialyzed iCP at a molar ratio of 10:1 and incubated at 37°C for 30 min. Then glutaraldehyde, at a final concentration of 0.1% (vol/vol), was added into this system, which was further incubated at 4°C for 2 h. Tris-HCl (pH 7.4) at a final concentration of 50 mM was then added to this system to terminate the glutaraldehdye- induced crosslinking reaction. Finally, the reconstituted sample was purified using a glycerol gradient (15-45% glycerol (wt/vol), 25 mM Tris-HCl (pH 7.4), 10 mM MgCl_2_, 1 mM DTT) and subjecting it to centrifugation for 16 h at 37,000 rpm. Fractions containing PA28αβ-iCP proteasomes were identified by performing a peptidase activity assay and negative-staining EM (Fig. S1).

### Peptidase activity assay

The activity of the proteasome was monitored by performing a peptidase activity assay as previously described (Leggett et al., 2005). To quickly determine which fraction or fractions contained proteasomes, we added Suc-LLVY-AMC to a 96-well plate for each sample. The 96-well plate was incubated at 37 °C and visualized by using a gel image system (Tanon-1600, Shanghai, China). We continuously measured the proteasome proteolytic activity by continuously monitoring the fluorescence of free AMC for 10 min using a multimode microplate reader (BioTek).

### Cryo-EM sample preparation and data collection

Holey carbon grids (Quantifoil R2/1, 200 mesh) were plasma treated using a Solarus plasma cleaner (Gatan). A volume of 2 µl of the sample was placed onto a grid, then flash frozen in liquid ethane using a Vitrobot Mark IV (Thermo Fisher). Data collection was performed using a Titan Krios transmission electron microscope (Thermo Fisher) operated at 300 kV and equipped with a Cs corrector. Images were collected by using a K2 Summit direct electron detector (Gatan) in super-resolution mode with a physical pixel size of 1.32 Å. Each movie was dose-fractioned into 38 frames. The exposure time was 7.6 s with 0.2 s for each frame, generating a total dose of ∼38 e^−^/Å^2^. All of the data were collected using the SerialEM software package (Mastronarde, 2005) with defocus values ranging from -1.5 to -2.8 µm.

### Cryo-EM 3D reconstruction

A total of 3,170 micrographs were used for the structure determinations. All images were aligned and summed using MotionCor2 (Zheng et al., 2017). Unless otherwise specified, single-particle analysis was mainly executed in RELION 3.0 (Zivanov et al., 2018). After CTF parameter determination using CTFFIND4 (Rohou and Grigorieff, 2015), particle auto-picking, manual particle checking, and reference-free 2D classification, 274,210 particles remained in the dataset. The initial model was a single-capped PA26-20S proteasome, derived from the previous crystal structure (PDB: 1Z7Q) (Forster et al., 2005), and low-pass filtered to 60 Å using EMAN 1.9 (Tang et al., 2007).

For the reconstruction, one round of 3D classification was carried out and resulted in extraction of 28% good PA28αβ-iCP particles and 60% good free iCP particles. Then another round of 3D classification was performed to further separate double-capped PA28αβ-iCP-PA28αβ and single-capped PA28αβ-iCP particles. An auto-refine procedure was performed in RELION for every of these three kinds of particles to generate their corresponding maps. Afterwards, particles were sorted by carrying out multiple rounds of 3D classifications, yielding a PA28αβ-iCP dataset with 45,030 particles, a PA28αβ-iCP-PA28αβ dataset with 24,627 particles, and a free iCP dataset with 56,663 particles. These particles were re-centered and polished, and one more round of auto-refine procedure was performed, resulting in a 4.1-Å-resolution map of PA28αβ-iCP, a 4.2-Å-resolution map of PA28αβ-iCP-PA28αβ, and a 3.3-Å-resolution map of free iCP. These maps were sharpened by applying corresponding negative B-factors, estimated by using an automated procedure in RELION 3.0. For each of the reconstructions, the resolution was accessed based on the gold-standard criterion of FSC=0.143, and the local resolution was estimated by using ResMap (Kucukelbir et al., 2014).

### Pseudo-atomic-model building

We used the X-ray crystal structures of the bovine cCP (PDB: 1IRU) (Unno et al., 2002) and mouse PA28αβ (PDB: 5MX5) (Huber and Groll, 2017) as template to build the homology models of the bovine iCP and human PA28αβ, respectively, through the SWISS-MODEL webserver (Arnold et al., 2006). The well-resolved side chain densities throughout our bovine free iCP map and the iCP portion within the map of PA28αβ-iCP enabled us to amend and refine the entire atomic model. This refinement was carried out first using COOT (Emsley et al., 2010), and then using the *phenix*.*real_space_refine* program in Phenix (Adams et al., 2010). The final atomic model was validated using *phenix*.*molprobity*. After determining the relative spatial arrangements of PA28αβ and iCP, we also built a pseudo-atomic model for the PA28αβ- iCP complex following the same procedure. In addition, C-terminal tails of PA28αβ, which were not built in the initial homolog model, were resolved in our PA28αβ-iCP complex map. So we added and refined these tails using COOT (Emsley and Cowtan, 2004) and Phenix according to the corresponding map density. The validation statistics for the atomic models are summarized in Table S1. Figures were generated with either UCSF Chimera or ChimeraX (Goddard et al., 2018; Pettersen et al., 2004), as well as PyMOL (http://www.pymol.org).

### Cross-linking/mass spectrometry analysis

The purified PA28αβ and iCP were buffer exchanged to Hepes buffer with 1% glycerol and incubated at 37°C for 30 min and then cross-linked by bis(sulfosuccinimidyl) suberate (BS3), with a final crosslinker concentration of 0.25 mM. 20 mM Tris-HCl was used to terminate the reaction after incubation on ice for 2 h. Cross-linked complexes were precipitated with cooled acetone and lyophilized. The resulting pellet was dissolved in 8 M urea, 100 mM Tris pH 8.5, followed by being subjected to TCEP reduction, iodoacetamide alkylation, and overnight trypsin (Promega) digestion. The digestion was quenched using 5% formic acid. The tryptic peptides were desalted using a MonoSpin C18 spin column (GL Science) and then separated within a home-packed C18 column (Aqua 3µm, 75 µm*15 cm, Phenomenex) in a Thermo EASY-nLC1200 liquid chromatography (LC) system by applying a 60- minute stepwise gradient of 5-100% buffer B (84% acetonitrile (ACN) in 0.1% formic acid). Peptides eluted from the LC column were directly electrosprayed into the mass spectrometer with a distal 2 kV spray voltage. Data-dependent tandem mass spectrometry (MS/MS) analysis was performed with a Q Exactive mass spectrometer (Thermo Fisher, San Jose, CA). Raw data were processed with pLink software (Yang et al., 2012) and Proteome Discoverer 2.2 xlinkx.

## ACKNOWLEDGMENTS

We are grateful to the staffs of the NCPSS Electron Microscopy facility, Database and Computing facility, Protein Expression and Purification facility, Mass Spectrometry facility for instrument support and technical assistance. This work was supported by grants from the NSFC-ISF 31861143028, National Basic Research Program of China (2017YFA0503503), the CAS Pilot Strategic Science and Technology Projects B (XDB08030201), the NSFC (31670754 and 31872714), the CAS Major Science and Technology Infrastructure Open Research Projects, and the CAS-Shanghai Science Research Center (CAS-SSRC-YH-2015-01, DSS-WXJZ- 2018-0002). Z.D. was supported by a National Postdoctoral Program for Innovative Talents (BX201700262), NSFC (31800623), and the China Postdoctoral Science Foundation (2017M621550).

## Author contributions

J.C., Y.W., and Y.C. designed the experiments. J.C. and Y.W. purified the proteins, performed functional analysis and collected the cryo-EM data. J.C., Z.D., and Y.W. performed data reconstruction. C.X and J.C. performed model building. C.P., J.C., and C.X. performed the XL-MS experiments. J.C., Z.D., and Y.C. analyzed the structure and wrote the manuscript.

---

**Fig. S1.**
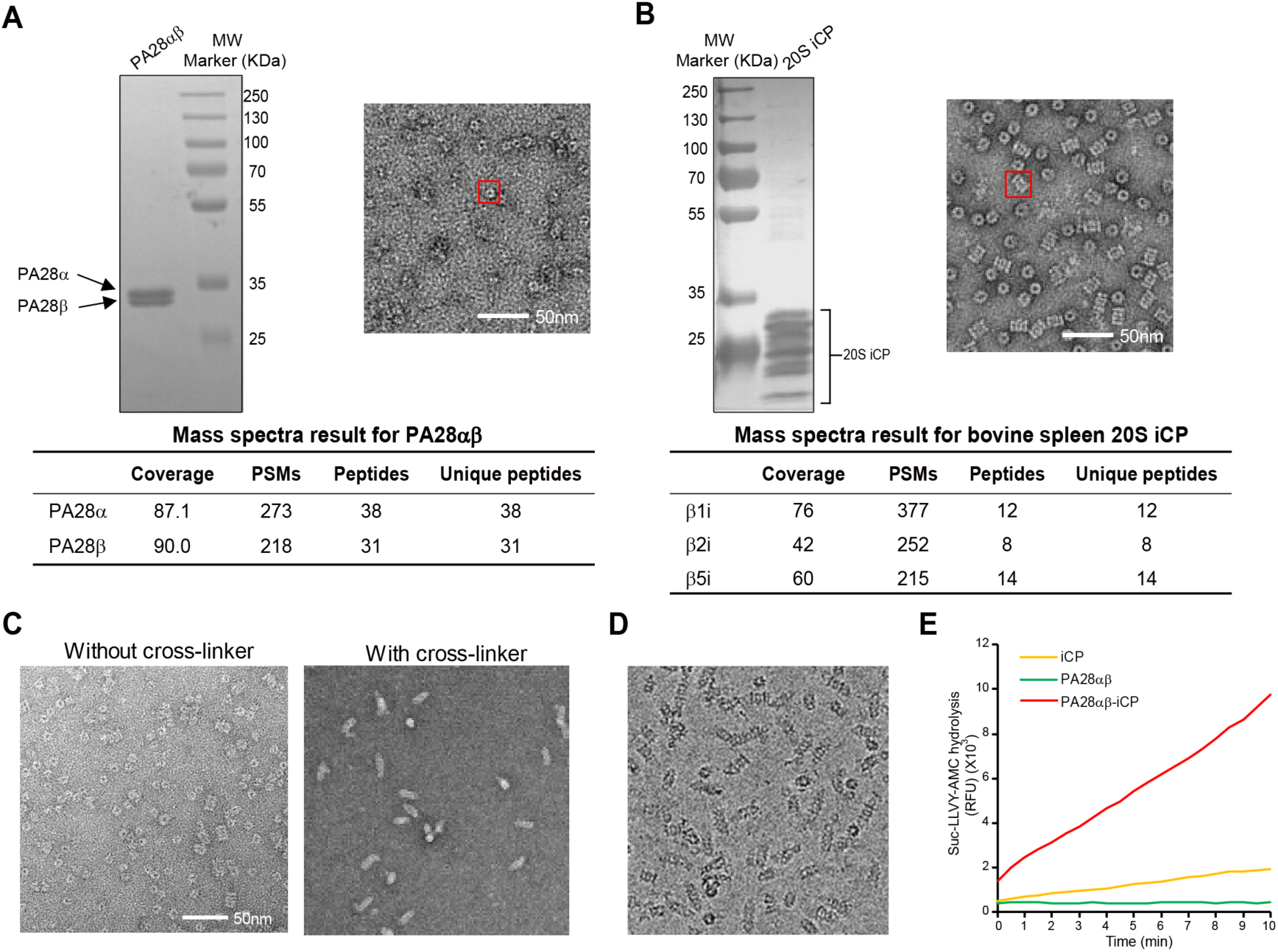
Purification and characterization of human PA28αβ and bovine spleen iCP. **(A-B)** Basic characterizations of **(A)** PA28αβ (human) and **(B)** iCP (bovine spleen) using SDS-PAGE, negative-staining electron microscopy (NS-EM), and mass spectroscopy analyses. PSMs stands for peptide spectrum matches, which were used to indicate the abundance of a certain protein. Unique peptides were used for protein identification. **(C)** NS-EM images of *in vitro* reconstitution of the PA28αβ-iCP proteasome with and without cross-linker. **(D)** A representative cryo-EM image of the *in vitro* reconstituted PA28αβ-iCP proteasome from purified PA28αβ and iCP in the presence of cross-linker. **(E)** Proteolytic activity assay of the reconstituted PA28αβ-iCP complex against the fluorogenic peptide Suc-LLVY-AMC. RFU, relative fluorescence units.

**Fig. S2.**
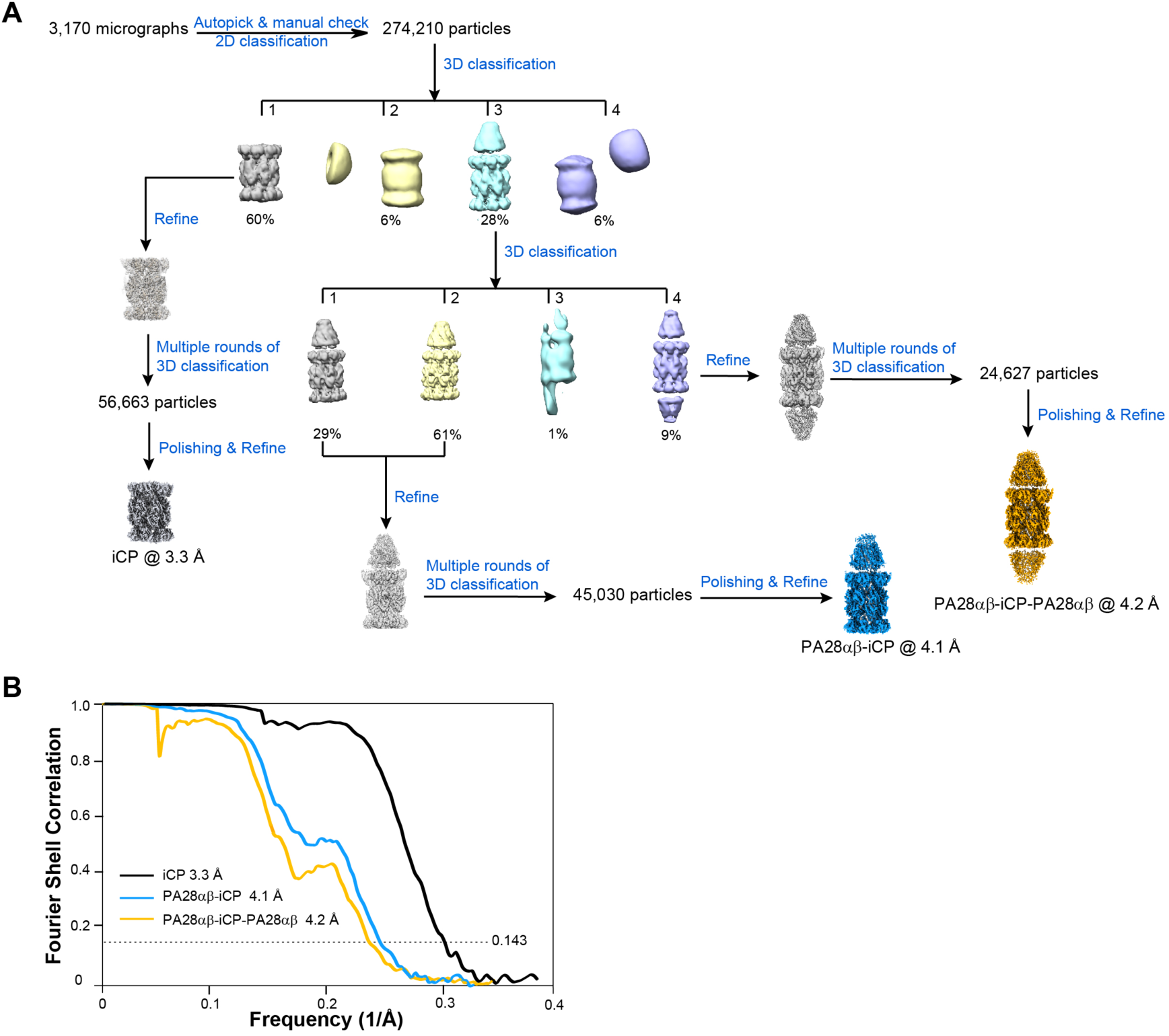
Structural analysis of the PA28αβ-iCP proteasome. **(A)** Work-flow of the PA28αβ-iCP proteasome data processing. More details are available in **Methods. (B)** Resolution assessment of our cryo-EM maps using Fourier shell correlation (FSC) at the 0.143 criterion.

**Fig. S3.**
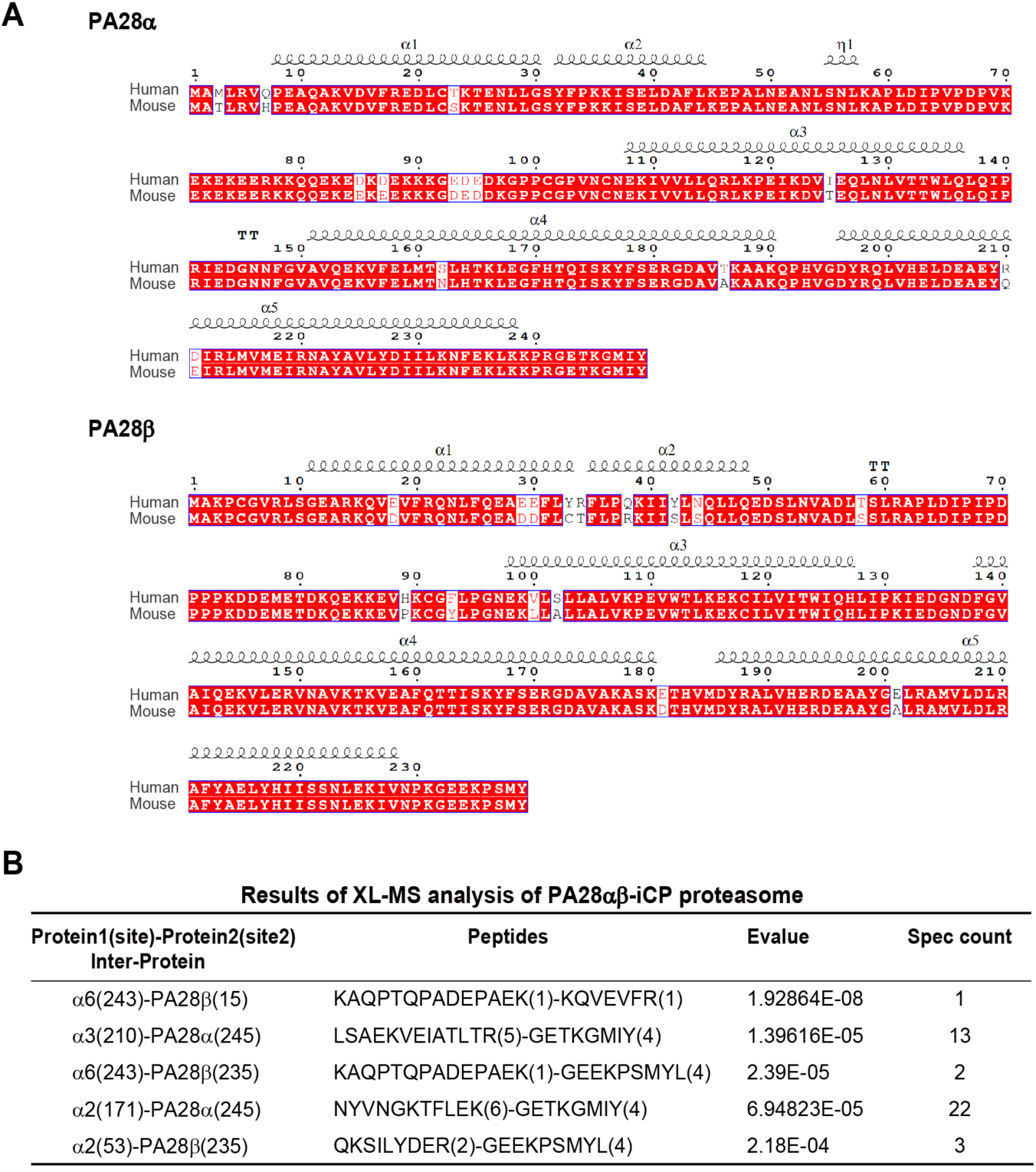
Sequence alignments of human and mouse PA28αβ and XL-MS results of PA28αβ-iCP proteasome. **(A)** Sequence alignments of human and mouse PA28α (top) and of human and mouse PA28β (bottom), generated by Espript. Showing sequence identities of 94.78% for PA28α between human and mouse, and 93.72% for PA28β between human and mouse. **(B)** Results of XL-MS analysis of the PA28αβ-iCP proteasome. Only the cross-linked contacts identified near the interface of PA28αβ-iCP are shown. We used an E-value of 1.00E^-02^ as the threshold to remove extra lower-confidence XL-MS data.

**Fig. S4.**
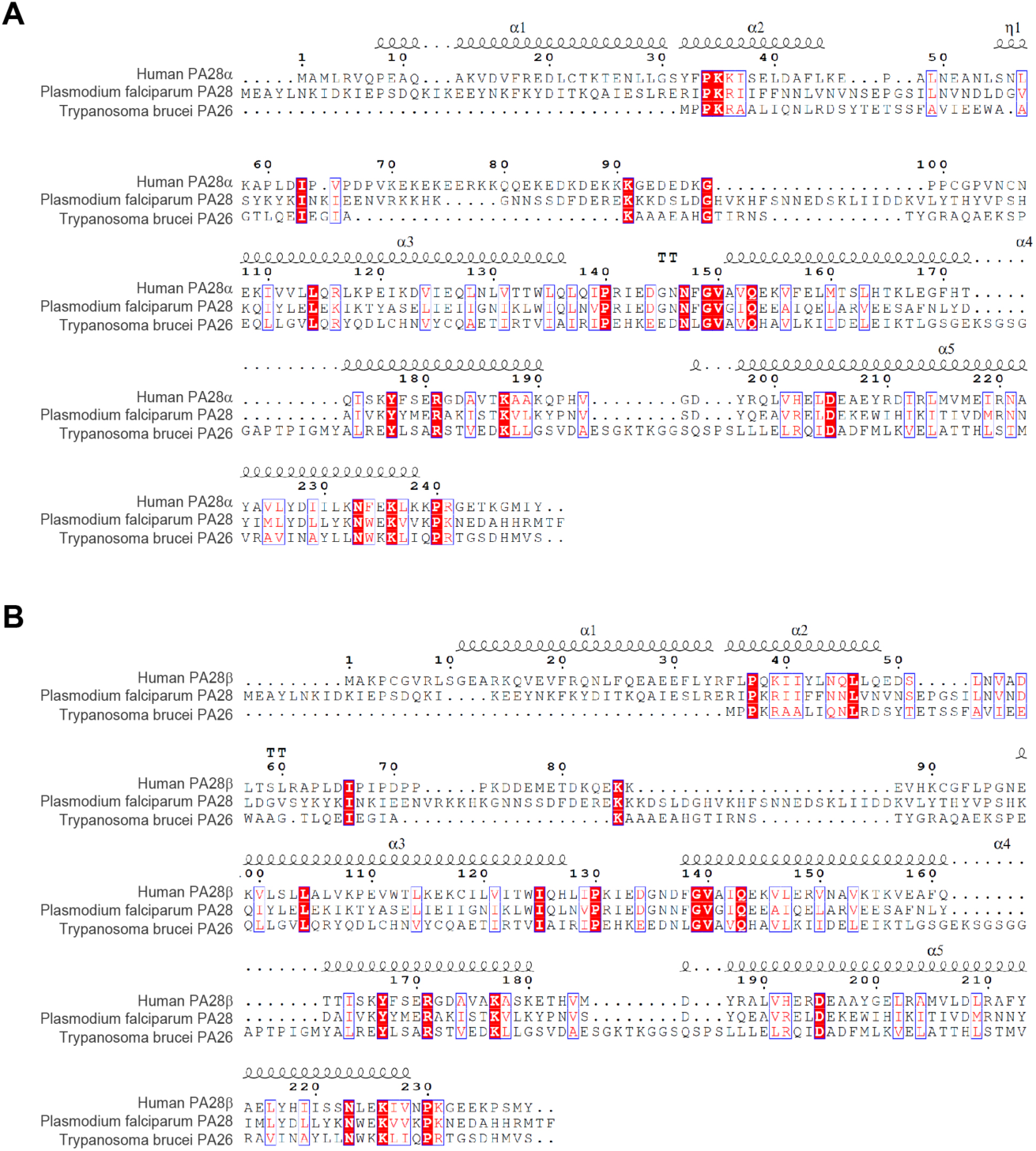
Sequence alignments of human, *Plasmodium falciparum* and *Trypanosoma brucei* 11S activator protein chains. **(A)** Espript representation of sequence alignment of human PA28α, *Pf*PA28 and *Tb*PA26, showing sequence identities of 26.76% between human PA28α and *Pf*PA28, and 15.64% between human PA28α and *Tb*PA26. **(B)** Espript representation of sequence alignment of human PA28β, *Pf*PA28 and *Tb*PA26, showing sequence identities of 24.38% between human PA28β and *Pf*PA28, and 12.46% between human PA28β and *Tb*PA26.

**Fig. S5.**
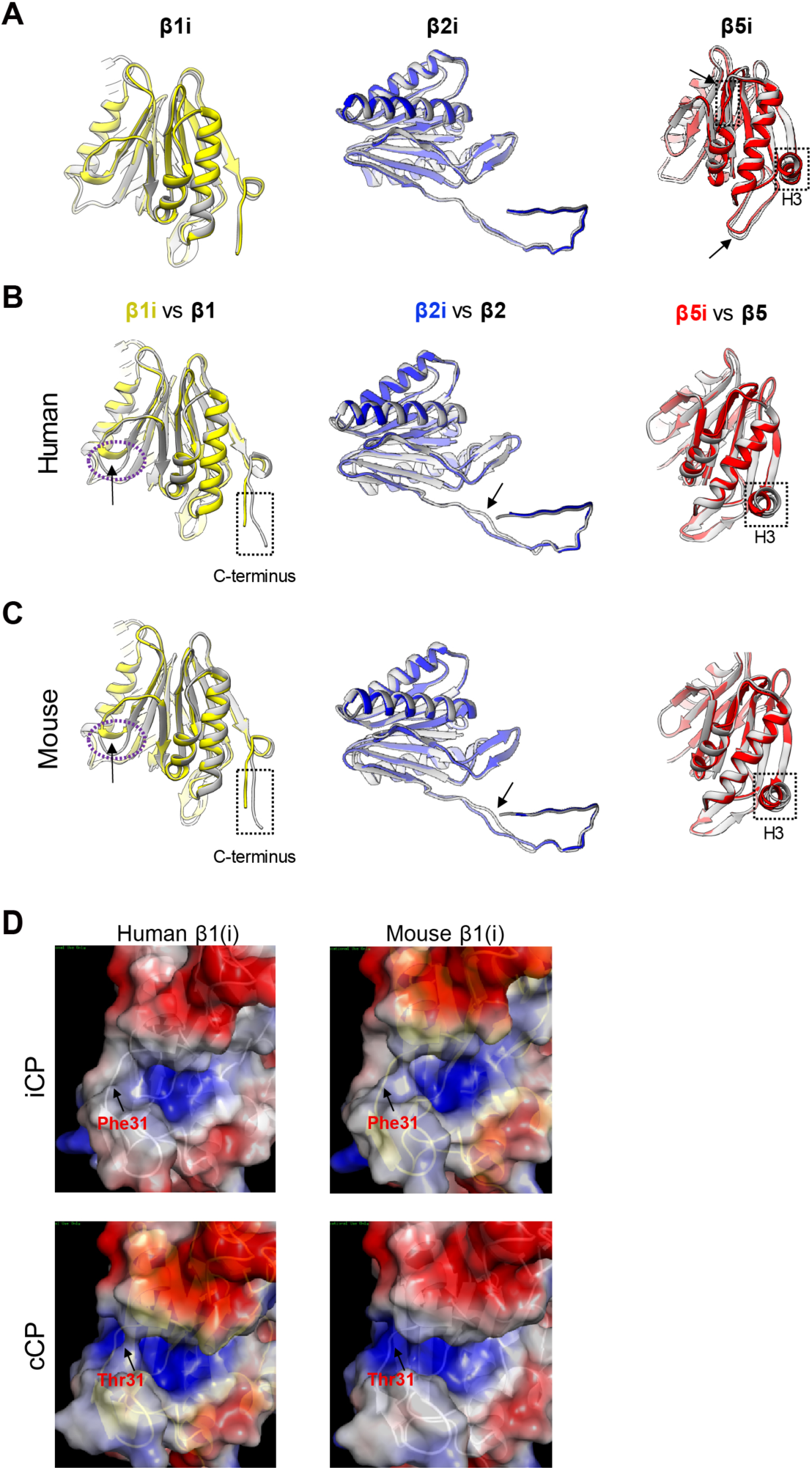
Structural comparison of the catalytic subunits between iCP and cCP. **(A)** Structural superpositions of the three catalytic subunits of our PA28αβ-bound iCP (with β1i, β2i, and β5i in color) on the corresponding subunits of our bovine free iCP (in gray). The observable conformational changes are indicated by black arrow and dotted frames. The rendering style is followed throughout. **(B-C)** Superpositions of the structures of **(B)** human iCP (PDB:6AVO) versus human cCP (PDB: 4R3O), and **(C)** mouse iCP (PDB: 3UNH) versus mouse cCP (PDB: 3UNE). **(D)** Surface charge representations of human β1i versus human β1, and mouse β1i versus mouse β1. With the most distinct residues between β1i and β1 in this region indicated.

**Table S1.** Statistics of cryo-EM data collection, processing, and model validation.

| PA28αβ-iCP |  |  |  |
| --- | --- | --- | --- |
| **Data collection** |  |  |  |
| EM equipment |  | FEI Titan Krios |  |
| Voltage (kV) |  | 300 |  |
| Detector |  | K2 |  |
| Pixel size (Å) |  | 1.32 |  |
| Electron dose (e<sup>-</sup>/Å<sup>2</sup>) |  | 38 |  |
| Exposure time (s) |  | 7.6 |  |
| Frames |  | 38 |  |
| Defocus range (μm) |  | -1.5~-2.8 |  |
| **Reconstruction** |  |  |  |
| Software |  | RELION 3.0 |  |
| Raw micrographs |  | 3,170 |  |
| Structures | PA28αβ-iCP | PA28αβ-iCP-PA28αβ | iCP |
| Final particles | 45,030 | 24,627 | 56,663 |
| Symmetry | C1 | C1 | C2 |
| Final overall resolution (Å) | 4.1 | 4.2 | 3.3 |
| Map-sharpening B factor (Å<sup>2</sup>) | -96 | -82 | -65 |
| **Refinement** |  |  |  |
| Software | Phenix & Coot | Phenix & Coot | Phenix & Coot |
| Rms deviations |  |  |  |
| Bond length (Å) | 0.0038 | 0.0106 | 0.0045 |
| Bond angle (°) | 0.88 | 1.08 | 0.95 |
| Ramachandran plot statistics (%) |  |  |  |
| Preferred | 94.97 | 97.50 | 96.12 |
| Allowed | 4.91 | 2.40 | 3.78 |
| Outlier | 0.12 | 0.10 | 0.10 |

